# Neural-vocal phase coupling reveals structured timing in birdsong production

**DOI:** 10.64898/2026.07.21.739824

**Authors:** Fiamma L. Leites, Bruno R. R. Boaretto, Cristina Masoller, Ana Amador

## Abstract

Understanding how neural population activity is temporally coordinated with behavior remains a central challenge in neuroscience. Songbirds provide a powerful model system for addressing this question because learned vocal production requires precise coordination among neural dynamics, temporally structured motor output, and auditory feedback. However, quantifying neural-vocal interactions is challenging because both neural and acoustic signals are rhythmic, noisy, and highly nonstationary.

Here, we investigate neural-vocal coordination during spontaneous canary singing using simultaneous recordings of neural population activity in a forebrain region of the song system and vocal behavior. Using a phase-resolved cross-correlation framework combined with surrogate-based statistical validation, we quantify neural-vocal interactions in short and highly variable song segments. Our analysis reveals that neural-vocal interactions are organized into distinct temporal regimes comprising positive, near-zero, and negative lags, consistent with neural activity preceding, accompanying, or following vocal output. The coexistence of these regimes is consistent with the integrative role of the recorded region, which receives auditory input, contributes to premotor control, and participates in the neural circuitry supporting song learning and the ongoing maintenance of adult song. We further find that correlated and anticorrelated interactions coexist throughout singing, with anticorrelated interactions consistently concentrated around near-zero lags. These anticorrelations identify periods in which decreases in neural population activity are closely aligned with sound production, revealing biologically relevant information that is obscured by analyses performed over complete song renditions.

Together, these results uncover a robust temporal structure linking neural population activity to vocal behavior and provide a broadly applicable framework for extracting transient neural-behavioral interactions from complex biological signals.

**Author summary:** Modern neuroscience can simultaneously record the activity of large neural populations, yet extracting meaningful relationships between neural activity and behavior remains challenging because natural behaviors are highly variable. Songbirds provide a unique opportunity to study this problem because their learned vocalizations share key features with human speech while remaining experimentally accessible. Here, we analyzed simultaneous recordings of neural population activity and vocal behavior in freely singing canaries. Instead of averaging neural activity across entire songs, we examined brief time windows and combined local correlation analysis with statistical tests to identify reliable interactions between the brain and behavior. We found that these interactions switch among several preferred timing patterns: neural activity can precede vocal output, occur at nearly the same time, or follow acoustic events. This diversity is consistent with the combined roles of the recorded brain region. Our work provides an intuitive and broadly applicable framework for uncovering transient neural-behavioral interactions that remain hidden by traditional time-averaged approaches. Because it requires only simultaneous recordings of two time series and makes minimal assumptions about their dynamics, the framework can be applied to many biological systems involving complex temporal signals.

## Introduction

Complex behaviors emerge from the coordinated dynamics of neural populations and the body across multiple temporal scales. Recent advances in large-scale neural recordings and population-level analyses have substantially improved our understanding of neural-motor interactions [1–4]. However, how neural population dynamics are temporally coordinated with behavior remains poorly understood [5–7]. This challenge is particularly significant when both neural and behavioral signals are rhythmic, noisy, and strongly nonstationary, making it difficult to identify meaningful temporal relationships between brain activity and behavior. Developing robust methods capable of extracting neural-behavioral interactions from such signals remains an important challenge in neuroscience.

Songbirds provide a powerful model system for investigating neural coordination during learned motor behaviors because they are one of the few animal groups capable of vocal learning, a rare ability shared with humans and a small number of other species [8, 9]. As a consequence, songbirds have become a valuable animal model for studying the neural mechanisms underlying learned vocal communication [10]. In addition, since learning and vocal production depend on the precise interaction between neural activity and temporally structured motor output, they are a powerful model to investigate neural-behavioral coordination [11–13]. Among songbirds, canaries (*Serinus canaria*) are especially well suited for studying neural-vocal timing relationships as their songs are composed of rhythmic sequences of repeated syllables spanning a broad range of production rates, from approximately 3 to 30 Hz [14–18]. This broad temporal repertoire generates diverse vocal rhythms that facilitate the investigation of neural-behavioral coordination across multiple timescales within a single behavioral context. These vocal dynamics emerge from activity within a specialized neural network known as the song system, in which the forebrain neural nucleus HVC plays a central role in song production and sensorimotor integration during singing [19–21]. Previous studies have revealed temporally structured activity in HVC during vocal production [22–26], supporting the idea that neural dynamics are closely linked to the temporal organization of song [27, 28]. However, characterizing neural-vocal coordination remains challenging because conventional analyses often rely on averages computed over extended time periods, potentially obscuring transient and heterogeneous temporal relationships.

More recently, computational approaches have increasingly focused on extracting temporally structured neural-behavioral interactions from high-dimensional and highly variable datasets, highlighting the need for analysis frameworks capable of resolving neural dynamics at behaviorally relevant temporal scales [6, 7, 29]. Recent computational methods have further emphasized the importance of phase-based and multivariate approaches for characterizing coordination between neural populations across multiple temporal scales [30, 31].

Different methodologies are available for quantifying the synchronization of two signals, *X*(*t*) and *Y* (*t*), and the results obtained can be qualitatively similar, or not, depending on the nature of the signals [32–34]. Because of its simplicity, cross-correlation analysis has been extensively used; however, caution is needed when assessing the statistical significance of cross-correlation values [35]. The cross-correlation function, *C*_*XY*_ (*τ* ), has been widely used to quantify the similarity between *X*(*t*) and *Y* (*t* − *τ* ) in signal analysis. For example, it has been used to analyze local field potentials recorded from different cortical regions [36]. The height and time shift of the main peak of *C*_*XY*_ (*τ* ), i.e., the peak that has the largest deviation from the background level, has been used to estimate the degree of synchronization and lag between *X*(*t*) and *Y* (*t*) [36]. Correlation-based approaches have also been used to study synchronization in the cat visual cortex, including trial-by-trial variations in neuronal response latencies and transient episodes of oscillatory synchrony captured through sliding-window cross-correlation and surrogate-based controls [37, 38]. Similarly, lagged correlations have been used to characterize how neural responses relate to speech dynamics in the human auditory cortex [39, 40]. This approach relies on the assumption that *C*_*XY*_ (*τ* ) has a well-defined, unambiguous peak, and that it is approximately symmetric, i.e., *C*_*XY*_ (*τ* ) ≈ *C*_*Y X*_ (−*τ* ). However, this assumption does not hold if *X*(*t*) and *Y* (*t*) are non-stationary, or if the time interval analyzed is not long enough. Moreover, the estimation of the degree of synchronization and lag between *X*(*t*) and *Y* (*t*) from the height and time shift of the main peak of *C*_*XY*_ (*τ* ) fails when the signals’ rhythmicity generates multiple peaks in *C*_*XY*_ (*τ* ), as occurs, for example, in music-like time series.

Alternative measures to quantify the synchronization of two signals include the phase locking value computed from the instantaneous phases of the signals extracted using the Hilbert transform [41]. However, the Hilbert technique only gives an accurate estimation of the phases if the signals are sufficiently narrow band and the duration of the analyzed time interval is sufficiently long. A related method based on coherence analysis [42] also requires that the signals are sufficiently narrow band. Statistical phase-locking indices have also been developed to detect transient synchronization episodes in oscillatory neural signals, even in noisy data; however, their reliability depends on the choice of surrogate data methods used for significance testing [43]. Another popular approach to infer correlations that reflect causal interactions is based on autoregressive models. As an example, the classic Granger causality quantifier [44] and its variants [45, 46] are popular tools for analyzing interactions between neuronal populations in brain circuits. However, autoregressive models have unknown hyper-parameters that have to be carefully selected [47], and can return spurious causal relationships when the signals analyzed are not stationary. A different class of methods relies on the analysis of the timing of events defined in the signals. Event coincidence analysis has been used in a variety of contexts to estimate the level of synchronization and temporal lag between two signals [48–51]. However, the results obtained with this approach can depend significantly on the criteria used to define the events and to decide whether the timing of two events is synchronous or not.

In this work, we develop a phase-resolved framework to investigate neural-vocal coordination during spontaneous canary singing. Our approach combines cross-correlation analysis with surrogate-based statistical validation and applies these tools to short sliding windows of simultaneously recorded neural and behavioral signals. Rather than relying on a single correlation estimate computed over an entire song, the method identifies statistically significant interactions at short temporal scales while preserving the temporal diversity of neural responses. Using this framework, we characterize the temporal organization of neural-vocal interactions during singing and show that neural-vocal coordination is not described by a single temporal relationship but instead involves multiple interaction regimes. More broadly, our results demonstrate the importance of temporally-resolved analyses for studying interactions between rhythmic and strongly nonstationary biological signals.

## Results

### Neural population activity exhibits structured phase relationships with vocal behavior

Figure 1 shows simultaneous recordings of HVC neural population activity and vocal production during spontaneous singing in canaries. The recordings reveal rhythmic fluctuations in both neural and vocal signals across multiple timescales. In many song segments, oscillatory patterns in the MUA appear to co-vary with fluctuations in the acoustic envelope, although the temporal relationship is not constant throughout the song. Instead, different song segments exhibit distinct relative timing patterns between neural activity and vocal behavior, suggesting that neural-vocal interactions may be dynamically organized across singing.

**Fig 1.**
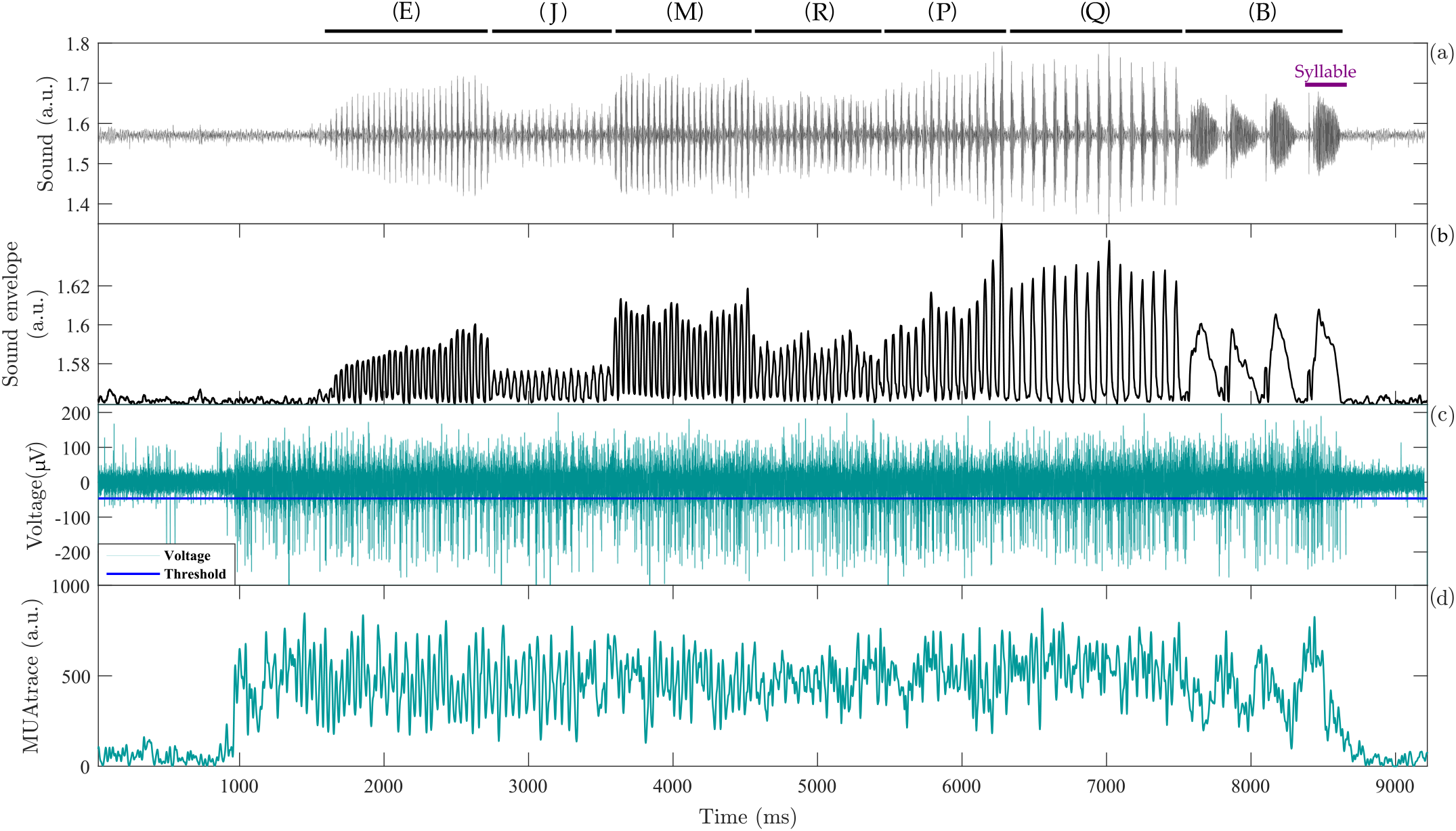
Simultaneous recordings of vocal behavior and neural population activity. a) Example oscillogram of canary song (Bird 1). Black bars indicate song phrases, and letters denote phrase identity within the bird’s repertoire. (b) Acoustic envelope of the song (ENV), highlighting the rhythmic temporal structure of vocal production. (c) Extracellular neural recordings obtained simultaneously from HVC during singing. (d) Multi-unit activity (MUA) derived from the extracellular recordings, providing a mesoscopic representation of coordinated neural population activity.

To characterize these neural-vocal interactions, we analyzed the cross-correlation functions between the MUA and ENV signals. Due to the rhythmic structure of birdsong, cross-correlation profiles frequently exhibited multiple peaks distributed across positive and negative lags, as illustrated in Fig. 2. The figure shows representative recordings from the three birds analyzed in this study. While both the neural and acoustic signals exhibit rhythmic structure, their corresponding cross-correlation functions contain several peaks of comparable amplitude rather than a single dominant maximum. As a result, identifying a unique temporal lag between neural activity and vocal behavior from the full-song cross-correlation is often ambiguous. These observations indicate that the temporal relationship between neural activity and vocal behavior cannot be adequately described by a single lag computed over the entire song. This complexity motivates the development of analytical approaches capable of resolving neural–vocal relationships at finer temporal scales and identifying dynamically changing lags throughout behavior.

**Fig 2.**
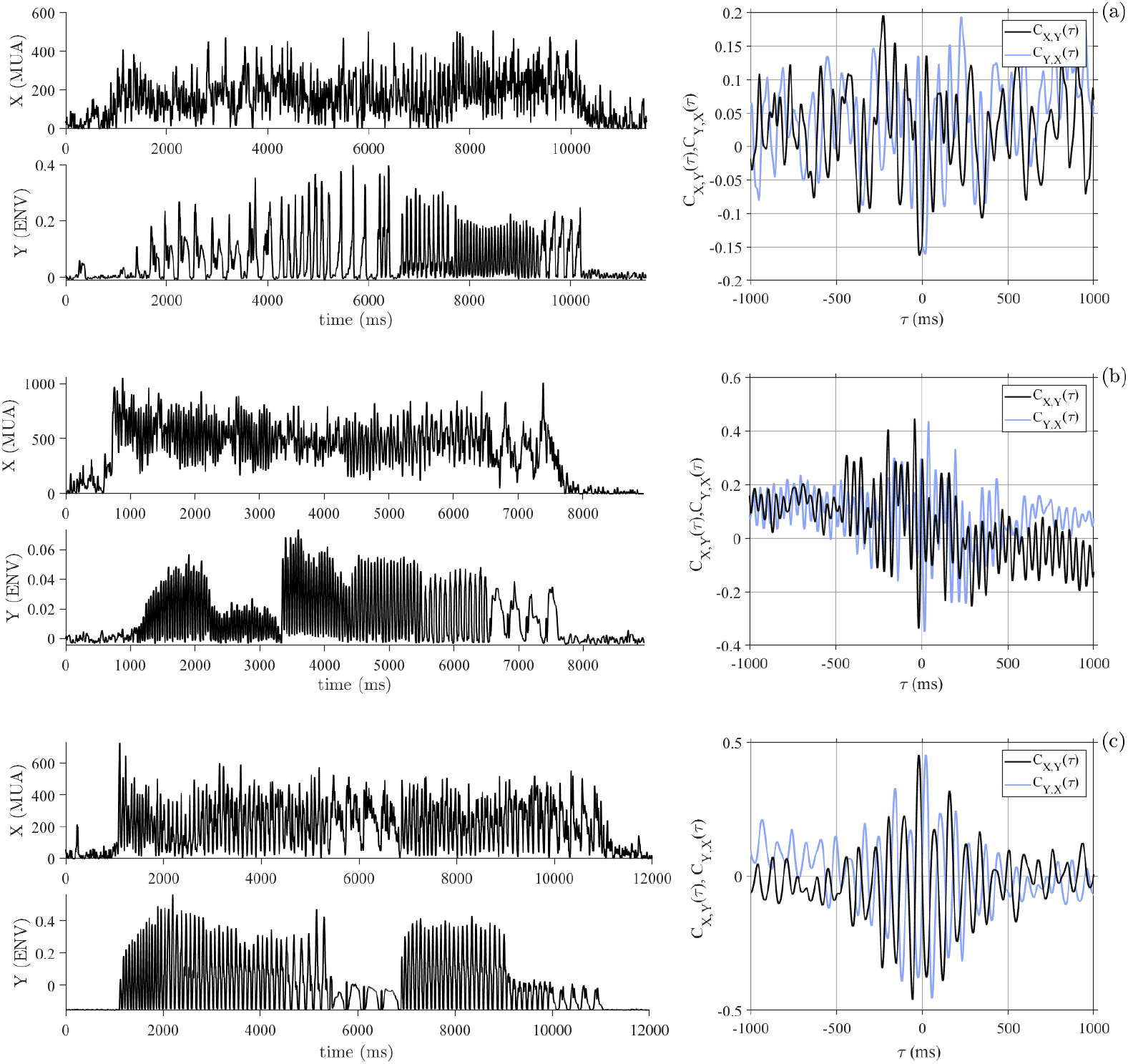
Examples of the time series of MUA (*X*) and ENV (*Y* ), and the corresponding cross-correlation functions, *C*_*XY*_ (*τ* ) and *C*_*Y X*_ (*τ* ). The analyzed interval corresponds to the full song duration (excluding the maximum lag, *τ*_max_ = 1000 ms, required for the cross-correlation computation). We show three representative datasets used throughout this study: (a) Bird 3, (b) Bird 2, and (c) Bird 1. These recordings serve as illustrative examples that are further developed in the following sections. Here, the cross-correlation is computed over the entire song, whereas later analyses apply a sliding-window approach to the same datasets.

### Phase-resolved analyses reveal temporal organization hidden by conventional correlation approaches

To investigate neural-vocal relationships at finer temporal scales, we computed cross-correlations in short sliding segments throughout the song. This approach allowed us to estimate both the cross-correlation value (*C*\*) and the corresponding temporal lag (*τ*\*) as a function of time. Positive values of *τ*\* indicate that fluctuations in MUA precede fluctuations in the acoustic envelope (ENV), whereas negative values indicate the opposite temporal ordering. For each segment, we computed the lagged cross-correlation functions *C*_*XY*_ (*τ* ) and *C*_*Y X*_ (*τ* ) between the MUA (*X*) and ENV (*Y* ) signals. Because rhythmic signals often generate multiple correlation peaks, particularly in short song segments, identifying a robust lag requires additional criteria beyond selecting the global maximum. We therefore retained only peaks that satisfied three conditions: they were sufficiently prominent, exceeded a minimum correlation magnitude (|*C*\*| > 0.5), and were statistically significant relative to surrogate and shuffled data (see Methods). In addition, candidate peaks were required to be approximately symmetric with respect to *τ* = 0 in the two cross-correlation functions. Figure 3 illustrates the procedure. In some segments, multiple peaks are present but none exceeds the statistical significance threshold, preventing the assignment of *C*\* and *τ*\* (Fig. 3a). In other segments, statistically significant peaks are detected, but they do not yield a unique lag estimate because *C*_*XY*_ (*τ* ) and *C*_*Y X*_ (−*τ* ) do not exhibit symmetric peaks (Fig. 3b,c). Only segments in which *C*_*XY*_ (*τ* ) and *C*_*Y X*_ (−*τ* ) contain statistically significant and approximately symmetric peaks were assigned values of cross-correlation and lag, *C*\* and *τ*\*, respectively (Fig. 3d). This procedure avoids identifying lags that arise solely from the rhythmic structure of the signals, providing a robust estimate of neural–vocal timing relationships.

**Fig 3.**
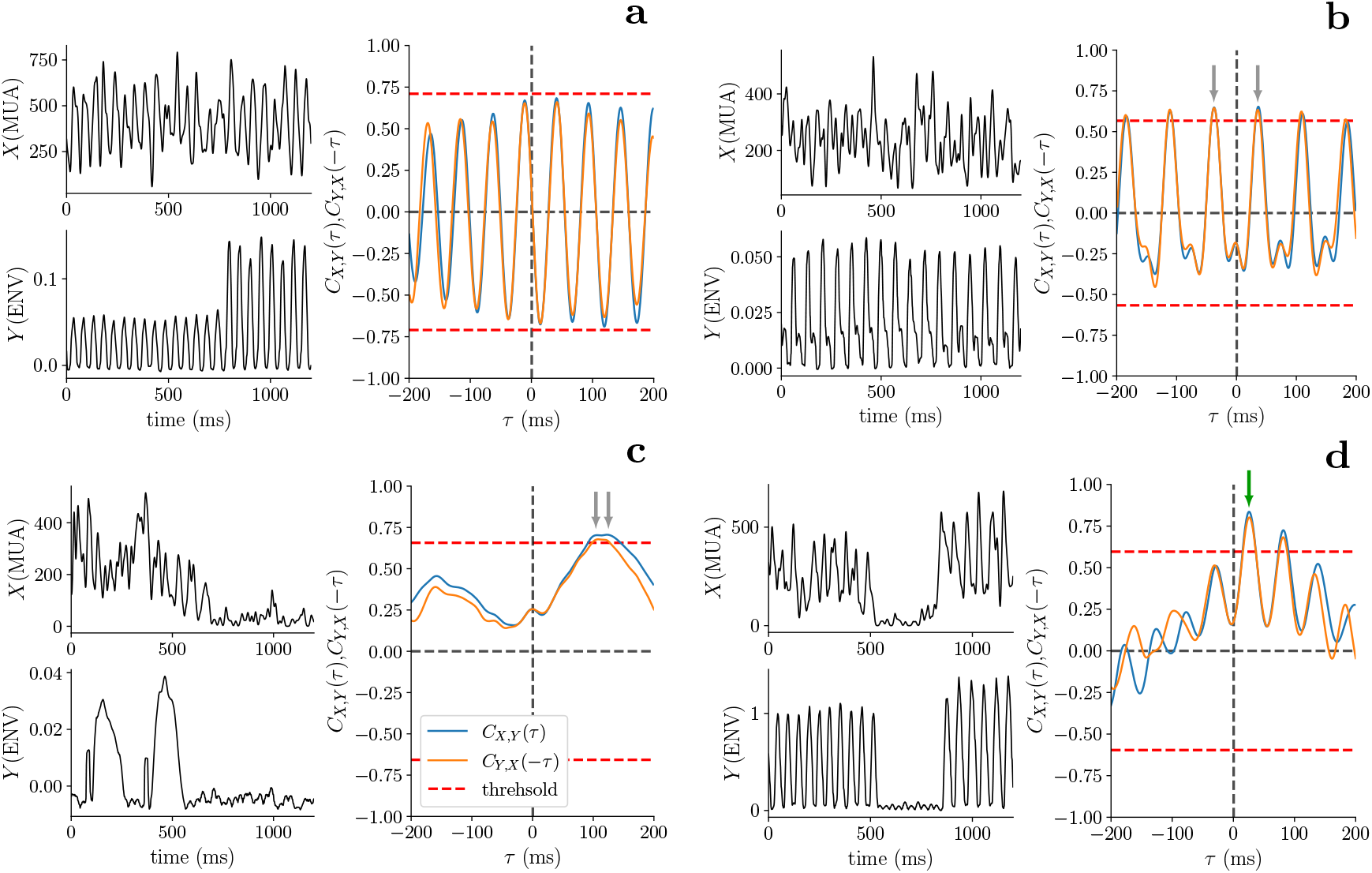
Examples of cross-correlation analysis. For each example, the top left panel shows the MUA(*X*) signal, the bottom left panel shows the corresponding ENV(*Y* ) signal, and the right panel displays the cross-correlation functions *C*_*XY*_ (*τ* ) (orange) and *C*_*Y X*_ ( −*τ* ) (blue), computed with *τ*_max_ = 200 ms. Dashed red horizontal lines indicate the significance thresholds obtained from surrogate data. Candidate peaks are marked by gray arrows, while matching peaks identified in both directions are highlighted in green. Peaks were considered valid only if they exceeded both surrogate thresholds and a fixed minimum correlation value of 0.5. Panel (a) shows several candidate peaks, but none exceed the surrogate thresholds, and therefore no significant correlation is identified. In panel (b), two prominent peaks with similar amplitudes are detected at opposite lags, preventing an unambiguous estimation of the lag. In panel (c), significant peaks are detected, but they are not symmetric between *C*_*XY*_ (*τ* ) and *C*_*Y X*_ (−*τ* ), resulting again in an indeterminate lag estimation. Finally, panel (d) shows a clear and significant peak that is symmetric across both directions, allowing an unambiguous determination of the correlation and lag values.

Applying this framework across complete songs revealed that neural–vocal interactions were dynamically organized throughout singing (Figs. 4, S2, and S3). Figure 4 shows the temporal evolution of the cross-correlation values *C*\* and the corresponding lags *τ*\* for one representative recording site. Significant interactions occurred only during restricted temporal intervals rather than continuously throughout the song. Moreover, both the sign and magnitude of *C*\*, as well as the corresponding lag values *τ*\*, varied across the song. These changes were often substantial, including transitions between correlated and anticorrelated interactions and shifts between positive and negative lags. However, the detected values did not fluctuate randomly from segment to segment. Instead, similar values of *C*\* and *τ*\* tended to persist over consecutive song segments, forming temporally localized interaction regimes. Rather than being uniformly distributed throughout the song, lag values were concentrated around a limited number of preferred values. Different lag regimes appeared at different moments during singing and often remained stable over multiple consecutive segments before transitioning to a different regime. This temporal organization suggests that neural-vocal interactions are structured by a small number of recurring timing relationships, a possibility that we investigate quantitatively in the following section. Panels (e) and (f) show representative examples of the cross-correlation analysis for two selected song segments indicated in panel (d). For each segment, the corresponding MUA and ENV windows are shown together with their cross-correlation functions. In both cases, statistically significant and approximately symmetric peaks are identified, allowing the assignment of *C*\* and *τ*\*. Figures S2 and S3 show analogous results from two other birds indicating that temporally localized interaction regimes are consistently observed across animals.

**Fig 4.**
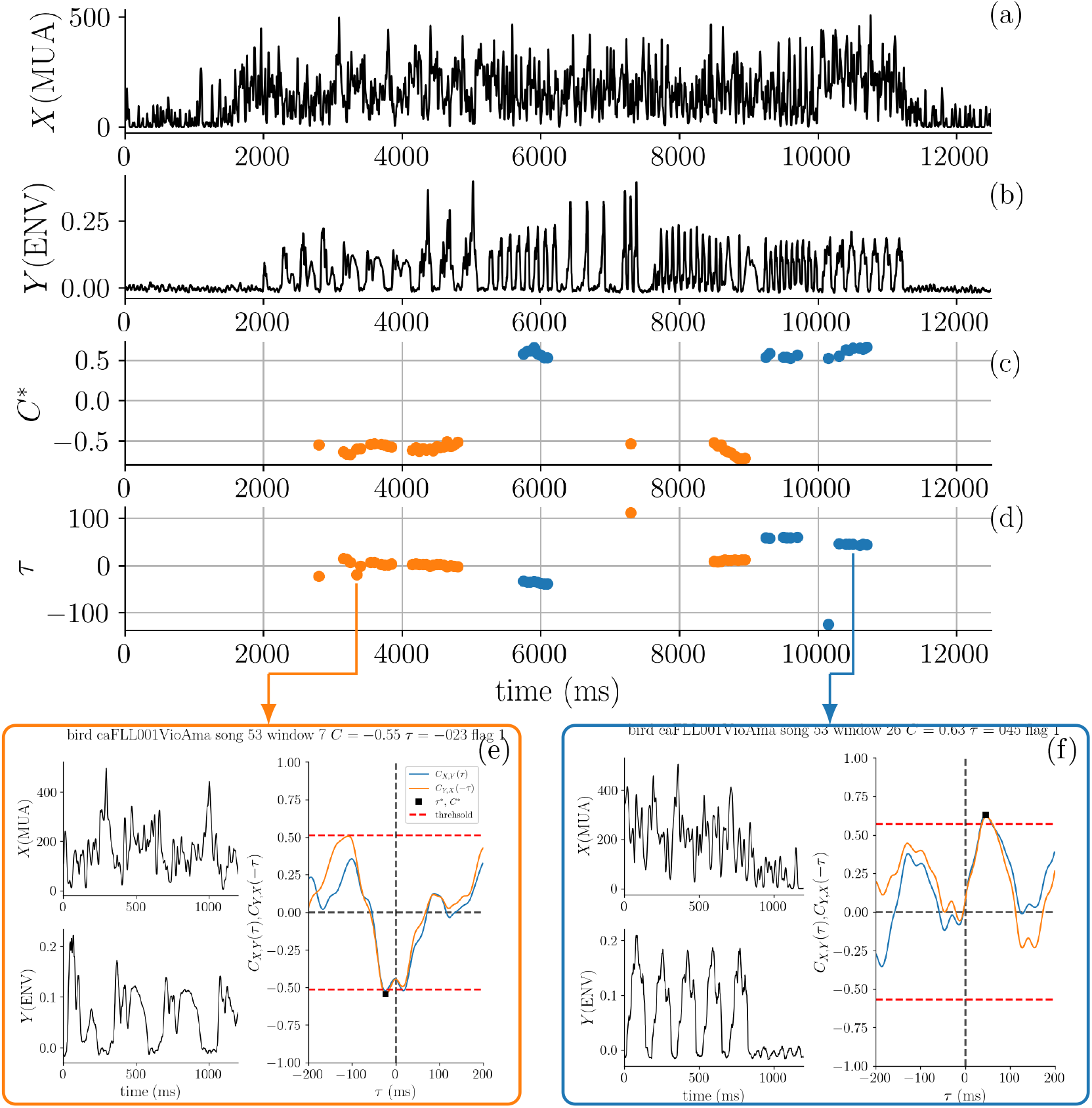
Temporal evolution of the cross-correlation values *C*\*and the corresponding lags *τ*\*. (a) *X* (MUA) and (b) *Y* (ENV) signals for Bird 3 (sample 837). Cross-correlations were computed using sliding windows of 1200 ms. Each point represents the correlation analysis of the subsequent 1200 ms segment beginning at that time point. (c) Cross-correlation peak values *C*\* and (d) corresponding lags *τ*\* computed as described in the Materials and methods section. Only statistically significant values according to both block-shuffled and surrogate-data tests are shown. Blue and orange dots indicate correlation and anticorrelation, respectively. The window step was reduced to 20 ms for visualization purposes. Panels (e) and (f) show representative examples of the cross-correlation analysis for two selected 1200 ms segments. In each example, the left panels show the MUA (*X*) and ENV (*Y* ) signals, while the right panel displays the cross-correlation functions *C*_*XY*_ (*τ* ) (orange) and *C*_*Y X*_ (−*τ* ) (blue), computed with *τ*_max_ = 200 ms. Dashed red horizontal lines indicate significance thresholds obtained from surrogate data, and statistically significant matching peaks are marked with asterisks.

### Neural-vocal interactions cluster into discrete temporal regimes

To determine whether neural-vocal interactions are associated with preferred temporal lags, we analyzed the distribution of *τ*\* values across birds and song segments. If lag values varied continuously, a broad distribution of *τ*\* values without clearly separated peaks would be expected. Instead, Fig. 5 reveals structured distributions with distinct peaks separated by regions of low probability density. To quantify this organization, we estimated the probability density of *τ*\* values using kernel density estimation and identified peaks and surrounding valleys as candidate temporal classes (see Methods). Across birds, the resulting distributions consistently revealed a small number of preferred lag values rather than a continuous distribution of lags. For example, the correlations of Birds 2 and 3 and the anticorrelations of all three birds exhibited a dominant class centered near *τ*\* ≈ 0, whereas the correlations of Birds 2 and 3 and the anticorrelations of Bird 1 also displayed multiple classes at distinct lag values spanning both positive and negative lags. Median values and interquartile ranges for all classes are reported in Table S1.

**Fig 5.**
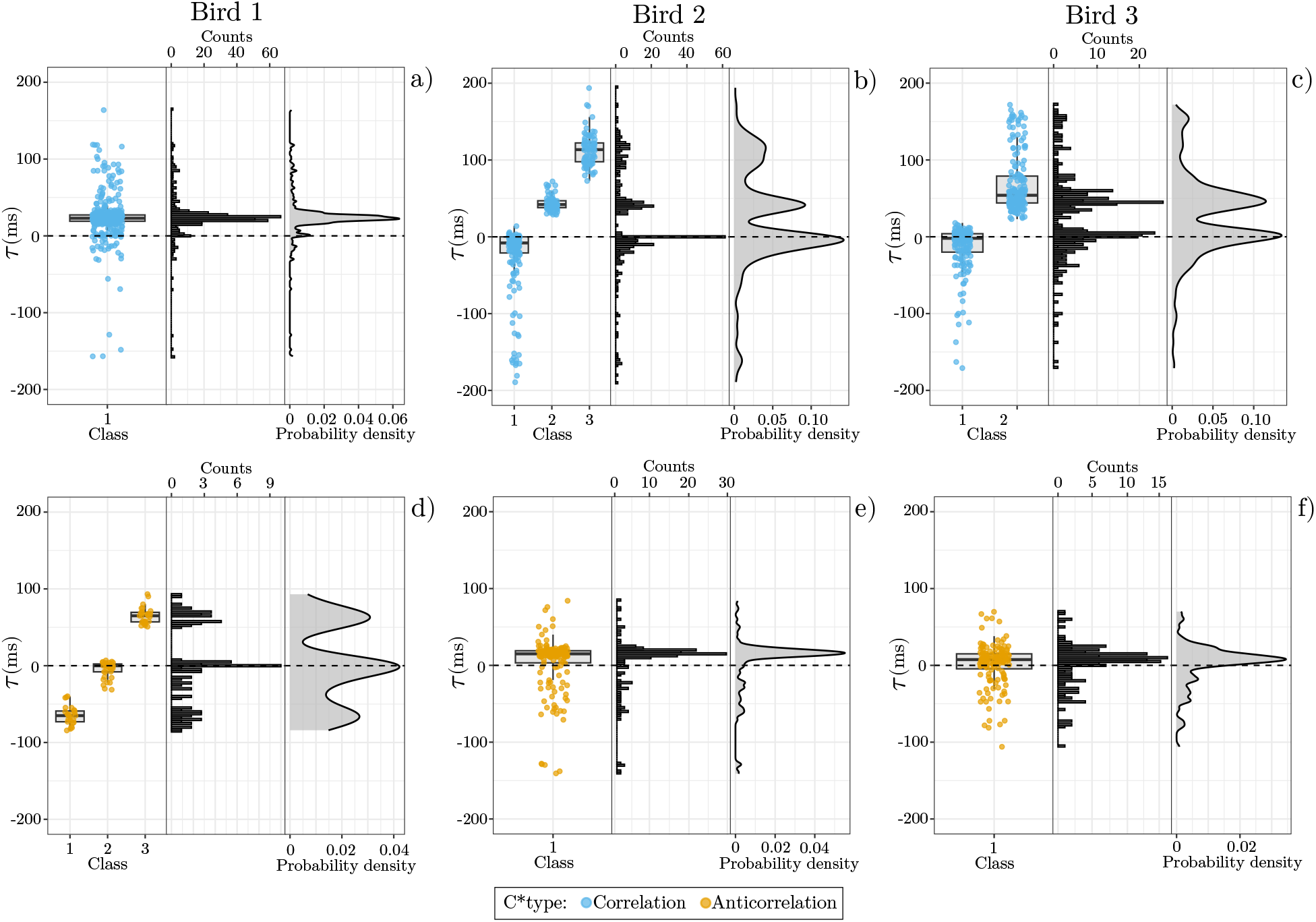
Neural-vocal interactions cluster into discrete temporal regimes. Each column corresponds to one bird dataset. Histograms and kernel density estimates of lag values (*τ*\*) reveal discrete temporal classes rather than continuous distributions. Boxplots summarize the phase regimes identified from the density peaks and surrounding valleys (see Methods). Blue and yellow points indicate correlated and anticorrelated interactions, respectively. Distinct phase regimes are consistently observed across birds, revealing structured temporal organization in neural-vocal coordination during singing. Detailed values in Table S1.

The distributions also exhibited extended tails toward both positive and negative lag values. In several datasets, lag magnitudes exceeded |*τ*\*| > 100 ms, indicating neural-vocal interactions operating over substantially longer timescales than those associated with the dominant classes. These large-lag events are visible both in the distributions of Fig. 5 and in the time-resolved analyses of Figs. S2, where they are associated with song segments containing slowly repeated or long-duration syllables. Although relatively infrequent, their recurrent presence across birds suggests that the temporal structure of vocal behavior can support neural-vocal coordination over a broad range of lags.

### Distinct phase regimes are associated with different neural-vocal interaction modes

To illustrate the temporal regimes identified above, we examined representative examples of neural-vocal interactions spanning positive, near-zero, and negative values of *τ*\* for both correlated and anticorrelated coupling (Fig. 6).

**Fig 6.**
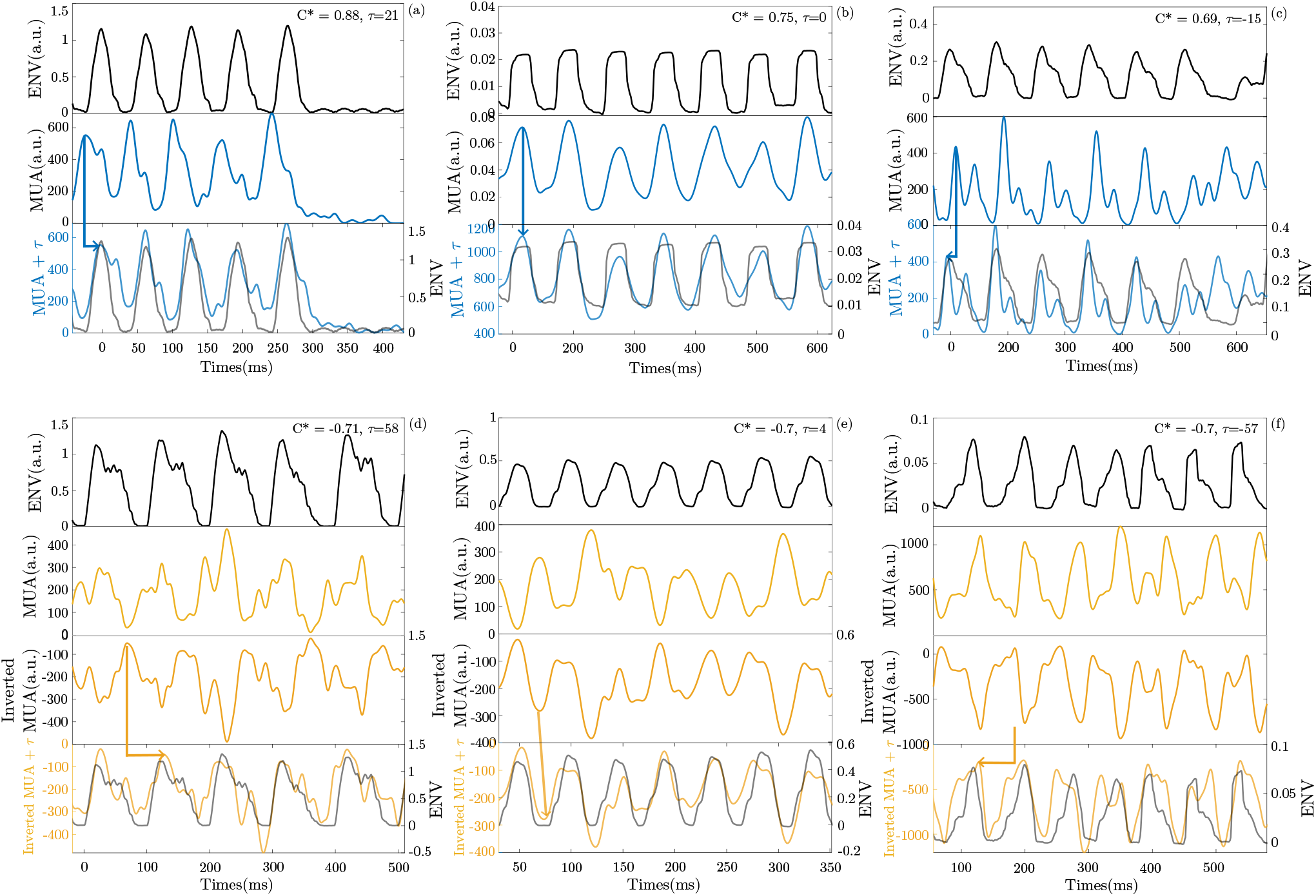
Representative examples of neural–vocal coupling regimes between the sound envelope (ENV) and the MUA. Each panel shows the envelope (top, black), the MUA (middle, blue or yellow), and their superposition after shifting the MUA by the estimated lag *τ*\* (bottom). In anticorrelation cases, the MUA is inverted prior to alignment to facilitate peak–trough matching. Arrows indicate the temporal shift applied to the neural signal. (a) Correlation (*C*\* > 0.5) with *τ*\* > 0: neural activity precedes the acoustic signal (premotor regime). (b) Correlation (*C*\* > 0.5) with *τ*\* ≈ 0: neural and acoustic signals are temporally aligned (synchronous coupling). (c) Correlation (*C*\* > 0.5) with *τ*\* < 0: neural activity follows the acoustic signal (sensory-driven regime). (d) Anticorrelation (*C*\* < −0.5) with *τ*\* > 0: neuronal suppression precedes the acoustic signal. (e) Anticorrelation (*C*\* < − 0.5) with *τ*\* 0: neuronal suppression is synchronized with sound production (antiphase). (f) Anticorrelation (*C*\* < −0.5) with *τ*\* < 0: neuronal suppression follows the acoustic signal.

For positively correlated interactions (*C*\* > 0), examples were observed in which neural activity preceded the acoustic envelope (*τ*\* > 0; Fig. 6a), was approximately synchronized with it (*τ*\* ≈ 0; Fig. 6b), or followed it (*τ*\* < 0; Fig. 6c). Analogous temporal relationships were also observed for anticorrelated interactions (*C*\* < 0), where decreases in neural activity were systematically associated with specific phases of the acoustic envelope (Fig. 6d-f). In each example, the lower panel shows the MUA signal shifted by the lag value *τ*\* estimated by the phase-resolved cross-correlation analysis. This alignment reveals the temporal relationship captured by the corresponding correlation peak and provides an intuitive visualization of the interaction regime. For correlated segments, the shift brings peaks and troughs of the neural and acoustic signals into alignment, whereas for anticorrelated segments the inverted MUA aligns with the corresponding acoustic fluctuations.

Together, these examples demonstrate that neural-vocal interactions encompass multiple coordination modes combining positive or negative correlations with positive, near-zero, or negative lags. They provide representative instances of the temporal regimes identified statistically in Fig. 5 and illustrate how the estimated lag values capture distinct patterns of neural-vocal coordination during singing.

## Discussion

Canary song provides a unique opportunity to investigate how neural population activity is coordinated with behavior across multiple timescales as learned vocal production generates rhythmic vocal patterns spanning a broad range of syllable repetition rates. Here we show that neural-vocal coordination is not characterized by a single temporal relationship, but instead is organized into multiple interaction regimes that appear during different portions of the song. A central result of this study is that the temporal lag between neural activity and vocal behavior is structured rather than continuously distributed. Within individual birds, *τ*\* values clustered around a small number of preferred lags, indicating that neural-vocal interactions are organized into discrete temporal regimes. These regimes included positive, near-zero, and negative lags, suggesting that different coordination patterns between neural activity and vocal behavior coexist during song production. Rather than remaining fixed throughout a song, these regimes appeared during specific song segments and persisted over short temporal intervals before transitioning to different interaction regimes.

The coexistence of positive, near-zero, and negative lag regimes reveals multiple modes of neural-vocal coordination. Positive lags are consistent with neural activity preceding vocal output, whereas negative lags are consistent with neural activity following acoustic events. Near-zero lags indicate approximately synchronous neural and vocal fluctuations. Although the present analysis does not establish causal relationships, the coexistence of these distinct interaction modes suggests that neural-vocal coordination during singing cannot be described by a single temporal mechanism. In addition to near-zero interactions, two birds exhibited prominent correlation classes centered at lags of approximately 40-60 ms. While the present analysis does not establish causal relationships, these timescales are compatible with the temporal lags expected between activity in HVC and the generation of vocal output through downstream motor pathways. The reproducibility of these lag classes across birds suggests that neural-vocal coordination may be organized around preferred temporal scales rather than continuously distributed lags.

Another prominent feature of the data was the recurrent presence of anticorrelated interactions. In particular, anticorrelated segments frequently clustered around *τ*\* ≈ 0 in all three birds. This observation suggests that neural-vocal coordination is not limited to periods in which neural and acoustic fluctuations vary in the same direction. Instead, coordinated interactions can also emerge through temporally structured anticorrelations. The consistency of this near-zero-lag regime across birds suggests that anticorrelated interactions constitute a robust component of neural-vocal coordination during singing.

We also observed interactions associated with larger lag magnitudes, in some cases exceeding |*τ*\*| > 100 ms. These events were typically associated with slowly repeated or long-duration syllables, suggesting that the temporal structure of vocal behavior influences the timescales over which neural and acoustic signals can become coordinated. Together, these findings indicate that neural-vocal interactions operate across multiple temporal scales during natural singing behavior, from near-synchronous coordination to lags spanning more than one hundred milliseconds.

An important aspect of the present framework is that it operates on mesoscopic neural population activity and does not require the identification of individual neurons. By combining MUA recordings with phase-resolved analysis, it becomes possible to characterize neural-vocal interactions throughout complete song renditions and across diverse vocal patterns. More generally, the approach developed here provides a way to identify temporally localized coordination regimes in rhythmic and strongly nonstationary signals. Beyond birdsong, this framework may prove useful for investigating how neural population activity is coordinated with complex natural behaviors across multiple timescales.

## Materials and methods

### Birds

We collected data from three adult male canaries (*Serinus canaria*), sourced from a local breeder. All recordings were performed at the University of Buenos Aires, Argentina during the spring-summer breeding season, a period of peak vocal activity in this species. Birds were housed individually in sound-attenuating boxes, maintained under a 14:10-hour light-dark cycle simulating summer daylight. Food and water were provided *ad libitum*. All procedures were reviewed and approved by the Institutional Animal Care and Use Committee of the University of Buenos Aires (CICUAL-FCEN, UBA). All efforts were made to minimize animal suffering.

### Surgical procedures

Surgeries were performed under isoflurane anesthesia (1–1.5% in 100% *O*_2_; Richmond) using a stereotaxic frame, with the bird positioned at a standard 45° angle between the ear bar axis and the beak. Feathers and skin over the skull were removed, and small openings were made in the outer bone layer to allow access for craniotomies and implant anchoring. Fine wires were inserted between skull layers to serve as support posts. The main craniotomy (∼ 500*µm* diameter) was made at coordinates +2.4 *mm* lateral and 0 *mm* anterior relative to the Y-point (the intersection of the midsagittal sinus and the transverse sinus, used as a stereotaxic reference in songbirds). After removing the dura, the electrode array was lowered into the brain using a hydraulic micromanipulator and protected with paraffin. A ground wire was placed in a secondary craniotomy in the contralateral hemisphere. The implant was secured using dental acrylic, which was applied between the skull layers and around the anchoring wires, forming a strong bond. The assembly was then sealed with cyanoacrylate and reinforced with a plastic protective cap.

### Electrode array and electrophysiological recordings

We recorded extracellular neural signals in the telencephalic nucleus HVC, a premotor area critical for song production and learning, during spontaneous singing, simultaneously with audio, following the protocol described in [52]. Each bird was housed alone but intermittently exposed to auditory and visual stimuli from other canaries to encourage vocalizations. Recording sessions continued as long as reliable neural signals and frequent singing were observed. Audio and neural signals were continuously monitored and automatically recorded when sound amplitude crossed a manually set threshold corresponding to vocal output. Electrophysiological signals were acquired using custom-built tetrode arrays made from low-impedance, polyimide-coated tungsten wires (12.7 *µ*m diameter, 99.95% purity; California Fine Wire), as per [53]. Tetrodes were mounted on a lightweight manual microdrive (a device that allows precise positioning of electrodes in the brain), adapted from the design described in [54]. Signals were amplified using an RHD2132 16-channel amplifier and digitized at 30 kS/s using the Intan Technologies RHD2000 USB interface. Audio was captured with a 20 Hz-20 kHz electret microphone connected to a MAX4466 amplifier board and fed into an analog input of the same interface.

### Data pre-processing

For each analysis, we used the recording of an entire song, including a few seconds of silence before its onset and after its offset to capture baseline neuronal activity. The raw neural signal was high-pass filtered using a third-order Butterworth filter with a 300 Hz cutoff to isolate spiking activity. Multi-unit activity (MUA) was obtained using an amplitude threshold set at three times the background noise level, estimated as *σ* = |*x*|*/*0.6745, where *x* is the high-pass filtered signal and *σ* is a robust estimator of the noise standard deviation [55]. To obtain the MUA, we applied a Gaussian kernel to generate smooth activity traces per channel (bandwidth = 5 ms, timestep = 0.033 ms). These parameters were chosen by visual inspection to capture temporal patterns of neuronal activity (periods of activity-silence) while minimizing noise across all phrase types, (repeated sequences of syllables — the minimal vocal units of canary song), considering their different temporal scales (2 to 27 Hz, reflecting the range of syllable repetition rates across phrase types). This smoothed curve was multiplied by the spike count to obtain an estimate of the absolute spiking rate, combining the temporal structure of the activity with the total number of detected spikes. Traces from channels of the same tetrode were averaged, yielding four MUA per song. The audio signal was processed computing a root-mean-square (RMS) envelope (ENV) using a sliding window of 500 samples, which captures the overall amplitude dynamics of the audio signal over time. MUA and ENV were downsampled to 1 kS/s to facilitate computational processing and reduce analysis time. All signals were pre-processed offline with MATLAB.

### Cross-correlation analysis

We analyzed the temporal relationship between MUA (referred to as *X*) and ENV (referred to as *Y* ) — representing neural activity and vocal output, respectively — in a segment of *N* data points, from time *t* to *t* + *N*, by computing the lagged cross-correlation functions,

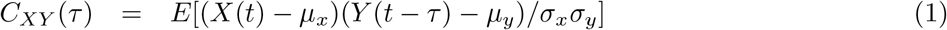

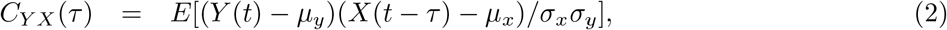

where *E* indicates the expected value and *µ*_*x*_, *µ*_*y*_, *σ*_*x*_ and *σ*_*y*_ are the mean and standard deviation of *X* and *Y* . If *X* and *Y* are stationary (i.e., their statistical properties do not change over time), *µ*_*x*_, *µ*_*y*_, *σ*_*x*_ and *σ*_*y*_ are constant in time and

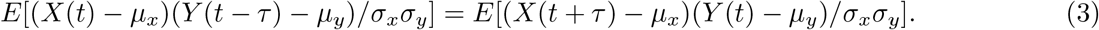

Therefore, *C*_*XY*_ (*τ* ) = *C*_*Y X*_ (−*τ* ). However, a visual inspection of the MUA and ENV signals shows that they are not stationary. Their mean values and standard deviations are not constant over time but show fluctuations that can be large when *N* is small. Therefore, in the segment [*t, t* + *N* ], *C*_*XY*_ (*τ* ) with *τ* in [−*τ*_max_, *τ*_max_] is estimated as:

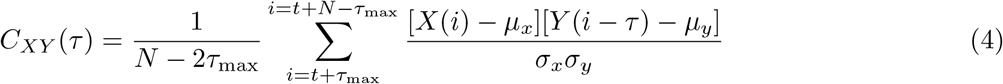

where *µ*_*X*_ and *σ*_*X*_ are the mean value and the standard deviation of *X*(*i*) with *i* in [*t* + *τ*_max_, *t* + *N* − *τ*_max_] and *µ*_*Y*_ and *σ*_*Y*_ are the mean value and the standard deviation of *Y* (*i*) with *i* in [*t* − *τ* + *τ*_max_, *t* − *τ* + *N* − *τ*_max_]. An equivalent expression is used to estimate *C*_*Y X*_ (*τ* ). We expect these estimates to be reasonable if *τ*_max_ << *N* [32]. Although the symmetry *C*_*XY*_ (*τ* ) = *C*_*Y X*_ (−*τ* ) does not hold for the nonstationary MUA and ENV signals, if the analyzed segment is long enough, *C*_*XY*_ (*τ* ) and *C*_*Y X*_ (*τ* ) have global maxima at 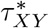 and 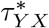 respectively, with 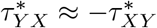 (see Fig. 2). This allows identifying the lag, *τ*\*, between the signals. We define 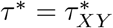, such that if *τ*\* > 0, fluctuations in MUA tend to precede fluctuations in ENV by *τ*\*, whereas if *τ*\* < 0, fluctuations in ENV tend to precede fluctuations in MUA by |*τ*\*|.

However, if the time window analyzed is short, the rhythmic structure of bird song, particularly pronounced in the ENV signal, leads to multiple peaks in *C*_*XY*_ (*τ* ) and *C*_*Y X*_ (*τ* ) (see Fig. 3), which make it difficult to identify a meaningful lag. The presence of multiple peaks in *C*_*XY*_ (*τ* ) and *C*_*Y X*_ (*τ* ) reflects the nontrivial temporal relationship between MUA and ENV. Therefore, we define the cross-correlation and lag, *C*\* and *τ*\*, between MUA and ENV in the segment [*t, t* + *N* ], as the height and lag of the peak that is 1) prominent, 2) significant (the significance tests are described in the next section), and 3) consistent, meaning that *C*_*XY*_ (*τ* ) and *C*_*Y X*_ (*τ* ) peak at 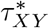 and 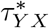 respectively, with 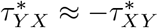. If several peaks satisfy these conditions, we choose the closest to *τ* = 0, while if there are no peaks that satisfy these conditions, *C*\* and *τ*\* in the segment [*t, t* + *N* ] are considered undefined. We note that, to capture both correlated and anticorrelated temporal relationships between MUA and ENV, the peak detection procedure is performed on the absolute value of the cross-correlation functions, |*C*_*XY*_ (*τ* )| and |*C*_*Y X*_ (*τ* )|. This allows identifying prominent and significant peaks regardless of their sign. Once a peak is selected, however, the corresponding non-absolute value of *C*_*XY*_ (*τ* ) or *C*_*Y X*_ (*τ* ) is retained, so that the sign of *C*\* indicates whether the interaction between the signals is predominantly correlated (*C*\* > 0) or anticorrelated (*C*\* < 0).

Specifically, the procedure to estimate *C*\* and *τ*\* is as follows:

1. For *X*(*i*) and *Y* (*i*) with *i* in the segment [*t, t* + *N* ], compute *C*_*XY*_ (*τ* ) and *C*_*Y X*_ (*τ* ) with *τ* in [−*τ*_max_, *τ*_max_] and *τ*_max_ ≪ *N* .
2. Identify candidate peaks in the absolute cross-correlation functions, |*C*_*XY*_ (*τ* )| and |*C*_*Y X*_ (*τ* )|. A peak is retained as a candidate only if it satisfies three criteria:
  a. *Prominence*: its prominence exceeds a threshold *p* = 0.1. This criterion excludes small local fluctuations arising from noise. The value *p* = 0.1 was selected because it provided a robust compromise between detecting genuine correlation peaks and rejecting minor fluctuations in the cross-correlation functions.
  b. *Correlation magnitude*: its absolute cross-correlation value exceeds a minimum threshold, |*C*\*| > 0.5, ensuring that only pairs of signals with a moderately strong temporal relationship are considered.
  c. *Statistical significance*: its magnitude exceeds the significance threshold estimated from surrogate and shuffled data (see Sec. IV F). Only peaks satisfying all three criteria are retained for further analysis.
3. Among the candidate peaks identified in the previous step, select the peaks of |*C*_*XY*_ (*τ* )| and |*C*_*Y X*_ (*τ* )| that are closest to *τ* = 0. We denote their heights and lags as 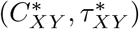 and 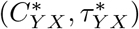, respectively. If no candidate peaks are found, *C*\* and *τ*\* are considered undefined.
4. Check consistency by verifying that 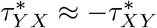. Specifically, we require that their difference is smaller than 10 ms (this value was selected after careful inspection of the peaks of |*C*_*XY*_ (*τ* )| and |*C*_*Y X*_ (*τ* )| across time windows of different durations). If this condition is not satisfied, *C*\* and *τ*\* are considered undefined.
5. When a pair of peaks is found that are prominent, significant, and consistent, they may have different heights. In such cases, we select the peak with the largest absolute value. If the selected peak belongs to *C*_*XY*_, then 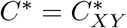 and 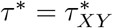; if it belongs to *C*_*Y X*_, then 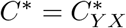 and 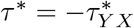. The sign of *C*\* is retained to distinguish correlated (*C*\* > 0) from anticorrelated (*C*\* < 0) interactions. If, however, the absolute heights of the two peaks differ by less than 10%, i.e., if 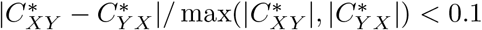, the result is considered indeterminate, as no clear criterion allows selecting one peak over the other.

The estimation of *C*\* and *τ*\* depends on a set of hyperparameters, namely the window size *N*, the maximum lag *τ*_max_, and the peak prominence threshold *p*. In this study, we set *N* = 1200. This choice reflects a trade-off between temporal resolution and statistical robustness: if *N* is too large, the window may include multiple distinct song phrases, thereby mixing different dynamical regimes and obscuring meaningful cross-correlations; if *N* is too small, the number of data points becomes insufficient to reliably estimate the cross-correlation, and the allowable range of lags is severely constrained. The maximum lag *τ*_max_ is chosen such that *τ*_max_ ≪ *N* . This is necessary because the estimation of *C*_*XY*_ (*τ* ) involves a loss of 2*τ*_max_ data points (see Eq. 4), reducing the effective sample size. If *τ*_max_ is too large, a substantial fraction of the data is discarded, compromising statistical reliability; if it is too small, the analysis cannot capture longer temporal lags between the signals.

The prominence threshold *p* determines the minimum prominence required for a local maximum to be considered a candidate peak. We set *p* = 0.1, which provided a robust compromise between detecting genuine correlation peaks and rejecting small local fluctuations in the cross-correlation functions. The additional threshold |*C*\*| > 0.5 ensured that only correlations strong enough to reflect a meaningful temporal relationship between the signals were retained for further analysis. We emphasize that these hyperparameters are dataset-dependent and should be adjusted according to the temporal structure, noise level, and sampling characteristics of the signals under analysis; there is no universal choice that is optimal across different datasets.

Figure 3 illustrates the peak extraction procedure. Each segment contains *N* = 1200 data points (1200 ms). The right panels show the cross-correlation functions *C*_*XY*_ (*τ* ) (blue) and *C*_*Y X*_ (−*τ* ) (orange) computed using *τ*_max_ = 200 ms. The red dashed line indicates the surrogate-based significance threshold. Panel (a) illustrates a segment in which several local maxima are present, but none exceeds the significance threshold. Consequently, no significant neural-vocal relationship is identified and *C*\* and *τ*\* are considered undefined for this segment. Panels (b) and (c) illustrate cases in which significant peaks are detected but do not yield a unique lag estimate. In panel (b), multiple candidate peaks are present, resulting in ambiguous lag assignment. In panel (c), peaks are detected in both cross-correlation functions but do not satisfy the symmetry criterion required for a consistent lag estimate. In both cases, *C*\* and *τ*\* are considered undefined. Panel (d) illustrates a segment in which a pair of statistically significant and approximately symmetric peaks is identified. In this case, a unique lag estimate is obtained and the corresponding values of *C*\* and *τ*\* are retained for further analysis.

### Statistical significance analysis

To assess the statistical significance of the cross-correlation magnitude and lag (*C*\* and *τ*\*) defined previously, we compare them against values obtained from two sets of surrogate datasets — artificially generated signals that serve as a null reference. *First surrogates dataset:* For each window of the MUA and ENV data, we generate 100 artificial samples using the Iterated Amplitude Adjusted Fourier Transform (IAAFT) method, implemented via a *Python* library [56]. The IAAFT procedure produces randomized versions of the original signals that preserve their frequency content and amplitude distribution, while destroying any temporal relationship between MUA and ENV. Figure S1 illustrates an example window of 1200 points from the MUA (a) and ENV (b) signals, extracted around 5.750 seconds from the data shown in Fig. 1(d) and Fig. 1(b), respectively. The panels below display different IAAFT surrogate samples generated from these original windows. For each pair of IAAFT windows, we compute the maximum of the cross-correlation in the same range of *τ* . In this case, for each window, we have a particular average *µ*_1_ and standard deviation *σ*_1_ computed over 100 IAAFT signals. *Second surrogates dataset:* Originally, the value of the cross-correlation of the signal is computed for a specific window *i*, extracted from the same time region for both MUA_*i*_ and ENV_*i*_. To evaluate whether the observed cross-correlation arises from the temporal structure within a window rather than from accidental similarities between unrelated segments, we compare it with the maximum cross-correlation value obtained from 1000 random pairs of windows, generated by independently selecting MUA_*j*_ and ENV_*k*_ windows at random, where *j* and *k* are indices within the same sample under analysis. In this case, each sample of the dataset produces a corresponding average *µ*_2_ and standard deviation *σ*_2_, which are common to all windows within that set.

### Statistical analysis and population segmentation of neural-vocal lags

To characterize the structure of neural-vocal lags, we analyzed the distribution of optimal lags *τ*\* obtained from the cross-correlation between the neural signal (MUA trace) and the acoustic envelope (ENV). Only segments with a well-defined *τ*\* were retained for further analysis. Observations with *C*\* > 0 were classified as correlated, whereas observations with *C*\* < 0 were classified as anticorrelated.

### Linear mixed-effects modeling

To evaluate whether *τ*\* differs between correlated and anticorrelated interactions, we fitted linear mixed-effects models assuming Gaussian errors and an identity link function:

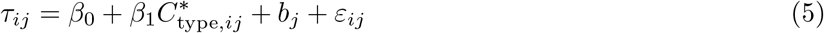

where *τ*_*ij*_ is the lag for observation *i* in bird *j*, 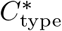 indicates correlation or anticorrelation, 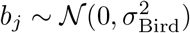 is a random intercept for each bird, accounting for baseline differences across individuals, and 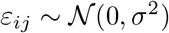 is the residual error capturing unexplained variability. The effect of correlation type was assessed using likelihood ratio tests by comparing the full model:

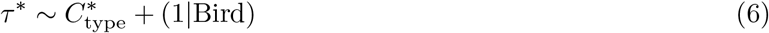

against a reduced model without this factor:

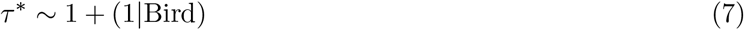

To evaluate whether recording site contributed additional explanatory power, we also considered a model including tetrode identity nested within bird:

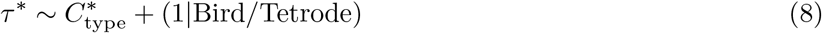

Model comparison was performed using likelihood ratio tests. Treating tetrodes as independent experimental units would constitute pseudoreplication, since recordings were obtained simultaneously within the same animal and brain region. Therefore, bird identity was considered the primary experimental unit for statistical inference.

### Within-bird analysis

To further assess the role of correlation type, analyses were also performed at the level of individual birds. Within each bird, models were compared with and without the 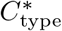 factor. This allowed evaluating whether the distinction between correlated and anticorrelated interactions explains systematic differences in *τ*\* at the level of individual birds.

### Class detection using kernel density estimation

To identify distinct regimes within the *τ*\* distributions, we implemented a data-driven segmentation based on kernel density estimation (KDE). For each analysis unit, the probability density function (PDF) of *τ*\* was estimated using a Gaussian kernel density estimator (KDE) with a fixed bandwidth adjustment factor of 0.65, which controls the degree of smoothing applied to the estimated density and was chosen to ensure consistent results across conditions. Local maxima (modes) of the density were detected and interpreted as candidate classes. Peaks were required to exceed a minimum prominence threshold of 1.5 times the median value of the estimated density function. Local minima (valleys) between adjacent modes were identified and used as population boundaries, requiring a density drop of at least 25% relative to the neighboring peak. Each *τ*\* value was assigned to a population based on the intervals defined by these boundaries. To avoid overinterpretation of sparse modes, only classes containing at least 10% of the total data points were retained.

### Statistical description

For each identified population, we computed robust summary statistics describing the corresponding *τ*\* distribution, including the median, first quartile (Q1), third quartile (Q3), interquartile range (IQR), and the number of observations.

## Supporting information

**Fig S1.**
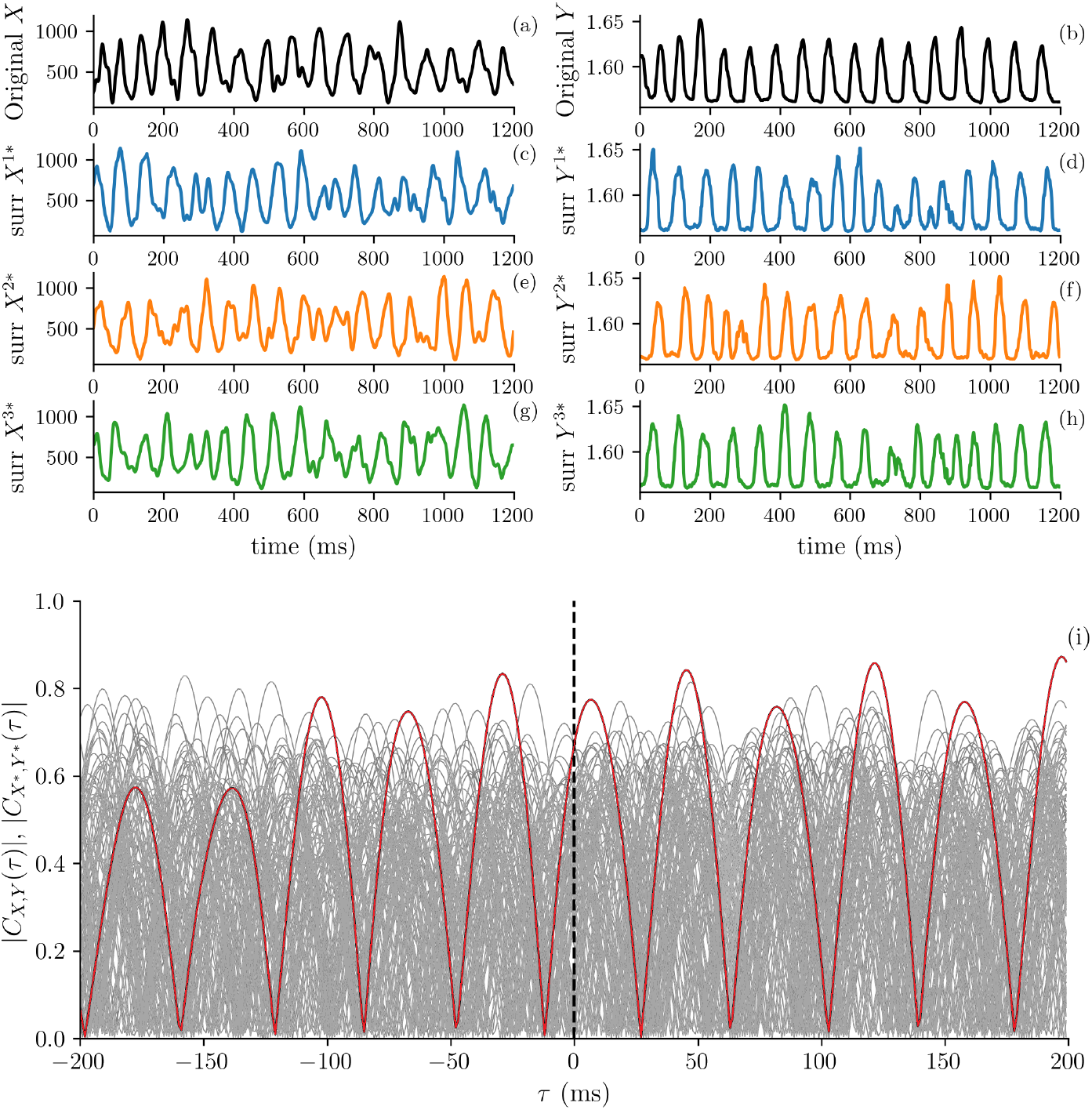
Example of Iterated Amplitude Adjusted Fourier Transform (IAAFT) surrogate generation. Panels (a) and (b) show 1200-point windows extracted from the MUA and ENV signals, respectively, centered around 5.750 seconds from the recordings in Fig. 1(d) and Fig. 1(b). The lower panels display representative IAAFT surrogate samples generated from the corresponding original windows. Panel (i) illustrates the cross-correlation function of the original signals (red solid line), along with those computed from 100 IAAFT surrogate samples (gray lines).

**Fig S2.**
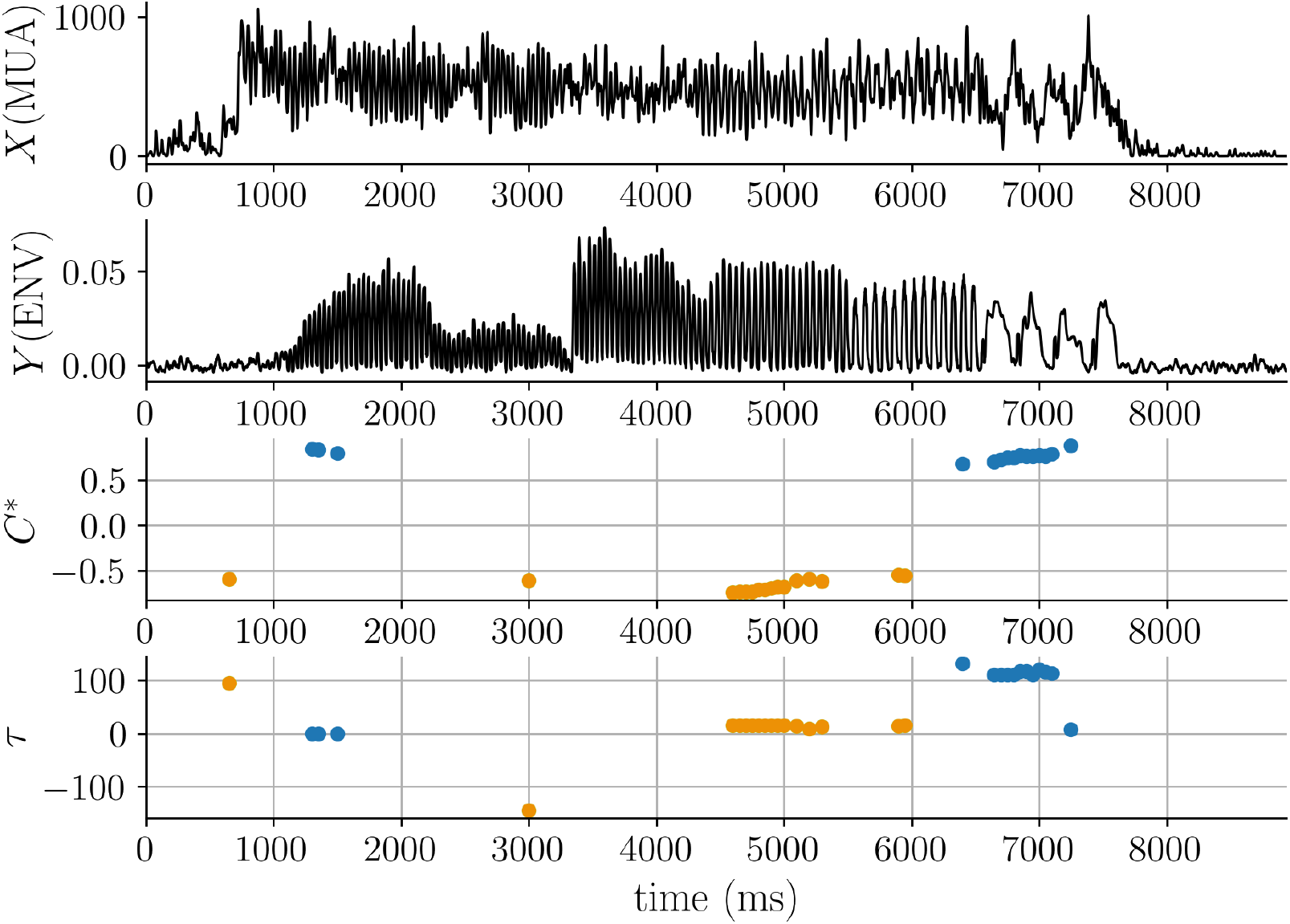
Temporal evolution of the cross-correlation values *C*\* and the corresponding lags *τ*\* for Bird 2 (sample 240) (a) *X* (MUA) and (b) *Y* (ENV) signals. Cross-correlations were computed using sliding windows of 1200 ms. Each point represents the correlation analysis of the subsequent 1200 ms segment beginning at that time point. (c) Cross-correlation peak values *C*\* and (d) corresponding lags *τ*\* computed as described in the Materials and methods section. Correlation values that were not statistically significant according to both block-shuffled and surrogate data tests are omitted. In (c) and (d), blue and orange dots indicate values where *C*\* is prominent, statistically significant, and satisfies the approximate symmetry condition *C*_*XY*_ (*τ* ) ≈ *C*_*Y X*_ (−*τ* ) within a small tolerance, corresponding to correlation and anticorrelation, respectively. The absence of points in (c) and (d) corresponds to windows where *τ*\* is undefined under the criteria of the method. The window step was reduced to 20 ms for visualization purposes.

**Fig S3.**
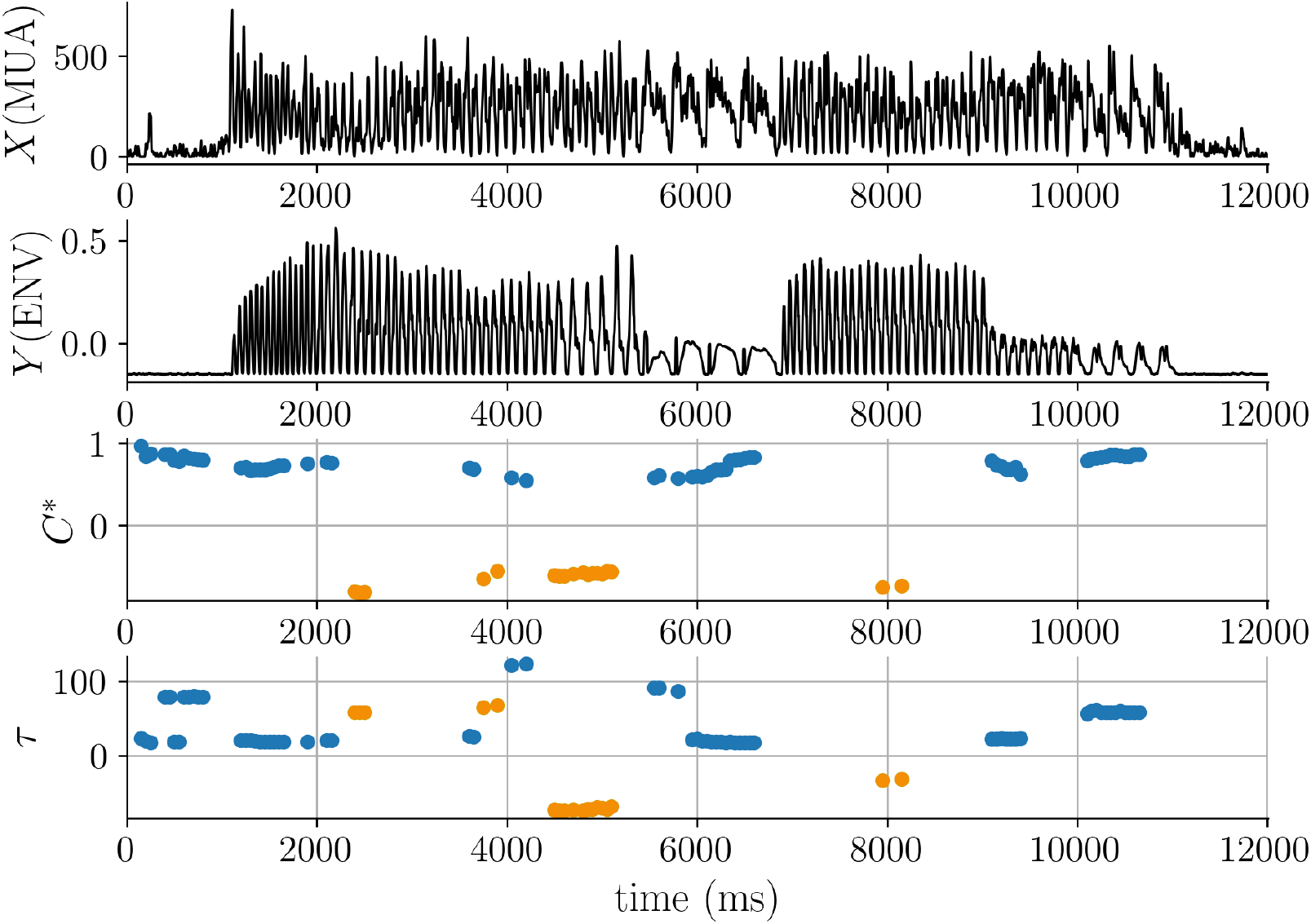
Temporal evolution of the cross-correlation values *C*\* and the corresponding lags *τ*\* for Bird 1 (sample 356) (a) *X* (MUA) and (b) *Y* (ENV) signals . Cross-correlations were computed using sliding windows of 1200 ms. Each point represents the correlation analysis of the subsequent 1200 ms segment beginning at that time point. (c) Cross-correlation peak values *C*\* and (d) corresponding lags *τ*\* computed as described in the Materials and methods section. Correlation values that were not statistically significant according to both block-shuffled and surrogate data tests are omitted. In (c) and (d), blue and orange dots indicate values where *C*\* is prominent, statistically significant, and satisfies the approximate symmetry condition *C*_*XY*_ (*τ* ) ≈ *C*_*Y X*_ ( −*τ* ) within a small tolerance, corresponding to correlation and anticorrelation, respectively. The absence of points in (c) and (d) corresponds to windows where *τ*\* is undefined under the criteria of the method. The window step was reduced to 20 ms for visualization purposes.

**Table S1.** Statistical summary of the *τ*\* distributions for each bird and group shown in Fig. 5.

| Bird | $C^*$ type | Class | $N$ | Median $\tau^*$ | Q1 | Q3 | IQR |
| --- | --- | --- | --- | --- | --- | --- | --- |
| Bird 1 | Correlation | 1 | 408 | 23 | 19 | 27.2 | 8.25 |
| Bird 2 | Correlation | 1 | 185 | -8 | -21 | 0 | 21 |
| Bird 2 | Correlation | 2 | 93 | 42 | 38 | 47 | 9 |
| Bird 2 | Correlation | 3 | 80 | 114 | 97.8 | 122 | 24.5 |
| Bird 3 | Correlation | 1 | 172 | -2 | -20 | 4 | 24 |
| Bird 3 | Correlation | 2 | 189 | 54 | 44 | 79 | 35 |
| Bird 1 | Anticorrelation | 1 | 25 | -65 | -73 | -59 | 14 |
| Bird 1 | Anticorrelation | 2 | 36 | 0 | -8 | 2.25 | 10.2 |
| Bird 1 | Anticorrelation | 3 | 27 | 65 | 57 | 69.5 | 12.5 |
| Bird 2 | Anticorrelation | 1 | 158 | 15 | 3.25 | 19 | 15.8 |
| Bird 3 | Anticorrelation | 1 | 148 | 7.5 | -4.5 | 15 | 19.5 |

Correlation corresponds to $C^* > 0.5$ , and anticorrelation corresponds to $C^* < -0.5$ . The table reports the number of valid $\tau^*$ values in each class ( $N$ ), the median $\tau^*$ , the first quartile (Q1), the third quartile (Q3), and the interquartile range (IQR). All $\tau^*$ values are expressed in milliseconds.

In Bird 1, correlation values form a single well-defined population centered at positive lags (median ≈ 23 ms). In contrast, anticorrelation values split into three distinct classes: one at large negative lags (median ≈ −65 ms), one near zero (median ≈ 0 ms), and one at large positive lags (median ≈ 65 ms).

In Bird 2, correlations distribute across three classes: one near zero lag (median ≈ −8 ms) and two at positive lags (medians ≈ 42 ms and ≈ 114 ms), none overlapping with those of Bird 1. Anticorrelations instead form a single population at small positive lags (median ≈ 15 ms), partially overlapping with the near-zero group in Bird 1.

In Bird 3, correlations organize into two classes: one near zero (median ≈ −2 ms) and another at positive lags (median ≈ 54 ms). The near-zero class partially overlaps with that of Bird 2, whereas no overlap is found with Bird 1. Anticorrelations form a single near-zero population (median ≈ 7.5 ms), overlapping with near-zero classes in both other birds.

